# Drug repositoning or target repositioning: a structural perspective of drug-target-indication relationship for available repurposed drugs

**DOI:** 10.1101/715094

**Authors:** Daniele Parisi, Melissa F. Adasme, Anastasia Sveshnikova, Yves Moreau, Michael Schroeder

## Abstract

Drug repositioning aims to find new indications for existing drugs, in order to reduce drug development cost and time. Currently, numerous successful stories of drug repositioning have been reported and many drugs are already available on the market. Although many of those cases are products of serendipitous findings, repositioning opportunities can be uncovered systematically by following either a disease-centric approach, as a result of a close relation between an old and new indication, a target-centric one, which links a known target and its established drug to a new indication, or a drug-centric approach, which connects a known drug to a new target and its associated indication. The three approaches differ in their complexity, potential, and limits, and most important the necessary starting information and computational power. Which one is predominant in current drug repositioning and what does this imply for future developments? To address these questions, we systematically evaluated over 100 drugs, 200 targets structures and over 300 indications from the Drug Repositioning Database. Each of the analysed cases has been classified based on one of the three repositioning approaches, showing that the majority, more than 60%, falls within the disease-centric definition, around 30% within the target-centric, and only less than 10% within the drug-centric. We therefore concluded that so far repositioning has mainly been disease and target repositioning and not, as drug repositioning, as expected by definition. We discuss the reasons and suggest directions to exploit the full potential of techniques useful for drug-centric in order to sustain future rationale repositioning pipelines.

## Introduction

### Drug repositioning to tackle pharmaceutical R&D decline

Drug discovery is a hard process, with an estimated success rate of only 2%^1^. Such a high chance of failures rises the average costs of patenting and selling a new molecule for pharmaceutical scopes to US $2–3 billion^2^. However, sometimes is possible to use approved drugs or molecules undergone phase trials to treat different conditions. A popular example is the case of sildenafil, meant to treat hypertension, successively commercialized against erectile dysfunction^3^. Also undesired effects can represent advantages towards different indications, such is the sadly known case of thalidomide, where the strong antiangiogenic activity became useful to treat multiple myeloma^3^. Investigating the efficacy of approved or discarded drugs for new indications with an approach called drug repositioning can in fact overcome some of the obstacles that are usually behind the failure of new molecular entity approach to the market, such as the necessity to meet the standard of quality set by previously marketed drugs^4^. Reducing the failure rate and therefore the average cost of the drug discovery process^5^, drug repositioning represents also a valid opportunity to find pharmacological tools against rare diseases^5,6^, and to make treatments of personilized medicine more affordable^7^.

### Drugs, targets, and diseases

Figure 1 shows a simplified classification of different repositioning approaches, where a protein target plays a key role in a disease, usually because of an alteration of its function, and a drug treats the disease by inhibiting or activating the target. Therefore, drug repositioning can focus on each of these three levels: disease, target, or drug. A focus on disease is the most direct approach, since it is driven by the hypothesis that a drug’s use can be expanded from the original to a closely related indication. As an example consider nilotinib, a tyrosine kinase inhibitor, approved for the treatment of imatinib-resistant chronic myelogenous leukemia^8^. A few years later, Novartis proposed the reposition of nilotinib to gastrointestinal stromal tumors. Disease-centric repositioning, as we define it, explores diseases of the same type, such as two types of cancer. The underlying assumption for disease-centric repositioning is that diseases of the same type have shared guiding principles. In the case of cancer this is e.g. summarized in the hallmarks of cancer^9^. However, despite such commonalities, indications differ and hence repositioning may not succeed. Actually, Novartis’ efforts to expand nilotinib to gastrointestinal stromal tumors were abandoned after a phase III trial concluded that it cannot be recommended^10^. Complementary to a disease-centric approach, target-centric repositioning builds on a novel link between a new indication and an established target. As an example, the protein tyrosine kinase ABL was recently suggested as novel player in Parkinson’s disease^11^, and therefore its inhibitors, such as nilotinib, may be effective against this syndrome^12^. This indication shift from cancer to neurodegeneration driven by the target ABL represents a case of target-centric repositioning. Drug-centric repositioning occurs when a novel target with its established link to a certain indication is predicted for the drug, as shown in Figure 1. For example, valproic acid is used in bipolar disorder and seizures because its ability to hit the mithocondrial enzymes Succinate-semialdehyde dehydrogenase (ALDH5A1) and 4-aminobutyrate aminotransferase (ABAT). However, due to its off-target interaction with the Histone deacetylase 2 (HDAC2), and the role of this protein in many types of cancers, it has been hypothesized to induce differentiation, growth arrest, and apoptosis in cancer cells, leading to the repositioning for the treatment of neoplastic conditions such as familial adenomatous polyposis^13^.

**Figure 1.**
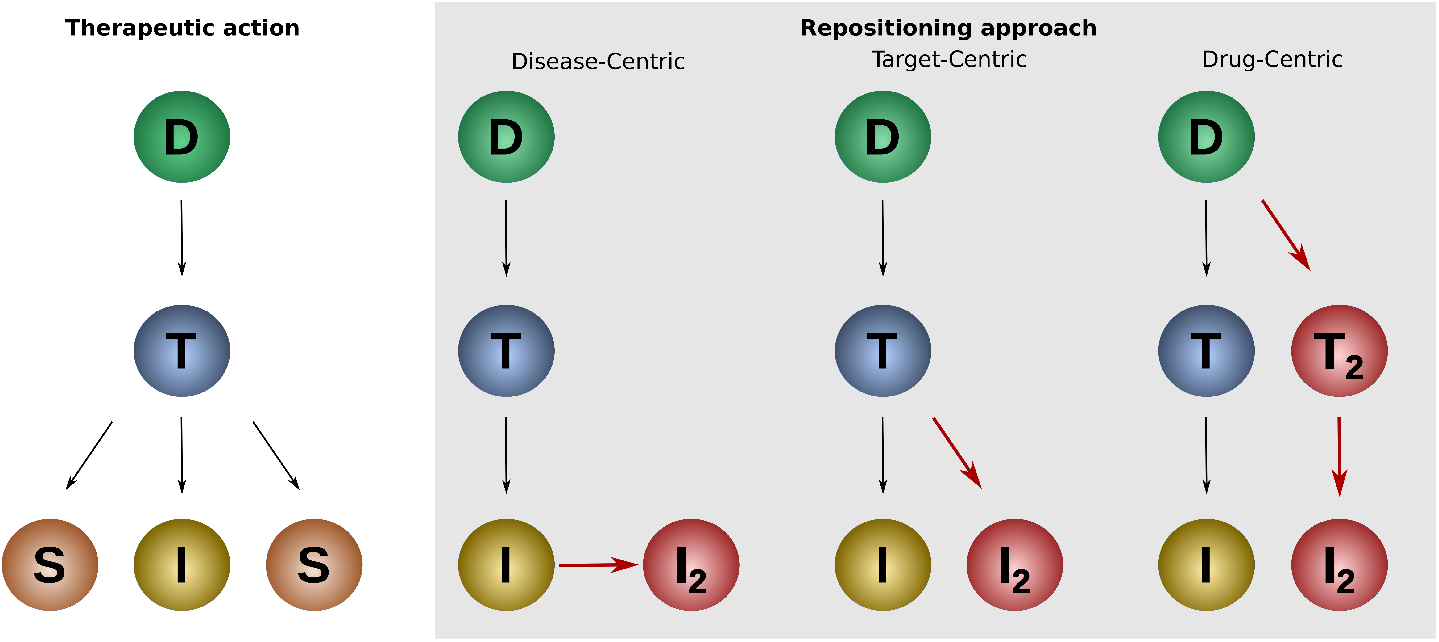
Different concepts behind drug repositioning. Relations among drug (D), targets (T) and siseases/indications(I) according to the different drug repositioning concepts. In disease-centric a drug’s application can be expanded from the original to a closely related indication, in target-centric the identification of a new indication is linked to a well established target and in drug-centric a new target is identified linking the drug to a new indication.

### Drug-target interaction prediction in drug repositioning

A precise identification of drug–target interactions allows users to compare different binding behaviours and therefore brings to the generation of novel rational repositioning hypotheses. Experimental identification of binding interactions can be challenging and expensive, therefore computational techniques for drug-target interaction prediction has gained a lot of attention. Computational approaches have been generally classified into ligand-based approaches, target-based approaches and machine learning-based methods^14^. With ligand-based approaches the binding is predicted by comparing the candidate ligand with compounds with known activity on the putative protein target. The performance of ligand-based approaches, such as QSAR and pharmacophore modeling, is related to the number of active ligands available for the protein target^15^. Target-based approaches, such as docking and binding-site similarity, are powerful tools for the identification of protein-ligand interactions based on the 3D structures of the target. Their limitations are related to the scarce availability of target structures, such in the case of GPCRs.^15,16^. Machine learning approaches predict novel drug-target pairs by using similarities among both compounds and targets. Those approaches are generally classified in feature vector-based machine learning and similarity-based machine learning. Similarity-based machine learning methods can be further grouped into three categories: kernel-based approaches, matrix factorization-based approaches and network-based approaches^17^. Compared with time consuming docking and information-demanding QSAR, machine learning methods can be faster and more efficient^18^. However, many limitations are related to the commonly used databases which contain only true-positive interactions and ignore many important aspects of the drug–target interactions, such as their dose-dependence and quantitative affinities.^19^.

### Structure based drug repositioning for new drug-target interactions

Several drug-centric approaches use structural information of the target active site or the complex drug-binding site to infer novel connections between drugs and targets. Many studies showed a correlation between drug-promiscuity and shared binding sites across the drug’s multiple targets, demonstrating the potential role of structural analyses of shared binding sites in drug repositioning^20^. Structure-based techniques, such as molecular docking, have been often applied on successful repositioning pipelines to predict new therapeutic candidates. For example, a docking-based approach was used to find novel targets for existing drugs by computationally screening the whole druggable proteome. As a result nilotinib was validated as potent inhibitor of MAPK14, adding potential to his role as antinflammatory drug^21^. A similar strategy was used to repurpose drugs against multi-drug resistant (MDR) and extensively drug resistent (XDR) tuberculosis. As a results, the anti-Parkinson drugs entacapone and tolcapone have been predicted to treat MDR and XDR tuberculosis since the structural and interaction similarity between the original target COMT and the new target InhA^22^. Docking scores have also been fused with other structural information with data integration techniques. For example, the method TMFS combined docking scores, ligand and receptor topology descriptor scores, and ligand-target interaction points, to predict potential new drug-target interactions and provide structural insight about their mechanism of action. In this way two novel drug-target interactions have been predicted and validated: mebendazole-VEGFR2 and celecoxib-CDH11^23^. Also several structure-based non-docking approaches found an extensive application in drug-repositioning in order to overcome the problem related with inefficiency and inaccuracy of docking. For example using information about the active-like state of the serotonin receptor 5-HT2C in complex with ergotamine and the inactive-like state of the same receptor in complex with ritanserin, was possible to suggest the extension of ergotamine pharmacological profile to the delta-opioid receptor^24^. Another non-docking structure-based approach used interactions patterns comparison to identify novel repositioning candidates against the cancer target Hsp27. Analysing the interaction patterns of the Hsp27 inhibitor brivudine was possible to indicate the approved anti malaria drug amodiquine as a promising anti-cancer agent^25^. Notwithstanding a rich history of successful cases, structure based drug repositioning suffers the limitation due to the scarce availability of structural information, concerning in particular certain classes of drug targets, such as GPCRs.

### Pros and cons of disease-, target-, and drug-centric repositioning

At first glance, disease-centric repositioning may appear faster and more direct than target- and drug-centric repositioning. In fact, a disease-centric repositioning hypothesis it’s based on a close connection drug-indication which might also avoid the deep knowledge about physicochemial interactions between drug and target. However, if this were the case, one cancer drug would cure all forms of cancer. Disease-centric approaches require a detailed understanding of the disease phenotype and underlying molecular processes to pursue the novel indication. Also, disease-centric approaches are hindered or supported (depending on the point of view) by patents. The drug under consideration and the old indication will be backed by patent claims, which tend to be broader than the old indication. Hence, the commercial exploration of a disease-centric repositioning needs to be closely coordinated with the relevant patent claims possibly limiting opportunities. Disease-centric approaches are suitable for systematic exploration. E.g. comprehensive, computational comparisons of phenotypes and drug side effects^26,27^ or comparisons of gene expression profiles^28^ define numeric similarities of diseases, which may drive a disease-centric approach.

In target-centric repositioning, we consider only drugs, for which old and new indication are of different type. Hence, it becomes less likely, that the new indication is already covered by patents of the drug. However, a novel link from target to new indication is a rare finding. And hence, these approaches are limited by the technology to uncover novel target-disease associations. Besides screening methods such as deep sequencing, micro-arrays, RNAi, which can give hints on candidate targets, the target-centric approach needs a deep molecular understanding of the relation of target and disease. The drug-centric pipelines are therefore the most indirect one, as the drug is only linked to the new indication via the discovery of a novel target, which is already established for the indication. Since each repositioning approach shows several pros and cons we performed a retrospective analysis to understand their distribution among the real successful drug-repositioning cases and the role of drug-target interaction prediction on indication switch of known drugs.

## Results

Which of the three approaches dominate drug repositonings? Is drug-target interaction prediction a driving force of drug repositioning? To address these questions, we analyzed all the repositioned small-molecule drugs active against a protein target, contained in the Repurposed Drug Database (RDD, http://www.drugrepurposingportal.com/repurposed-drug-database.php).

### Current Drug repositioning set contains 196 known cases

The merging of RDD with the Molecular Drug Targets (MDT) data led to a compiled report of 196 drug repositioning cases, 263 unique targets and 333 unique indications). Finally, 128 cases (194 merged cases excluding the cases regarding non small molecules and non protein targets) represents the starting point for our classification of repositioning cases (see Annex I).

### The majority of cases (59%) were Disease-centric discovered

In order to characterize the diseases susceptible to drug repositioning, we evaluated the frequency of appearance of the diseases in RDD for small molecule drugs by their root MeSH term key (results displayed in Figure 2). MeSH, Medical Subject Headings, is a comprehensive controlled vocabulary that provides a consistent way to retrieve information that may use different terminology to describe the same concept, facilitating indexing and searching. The most common MeSH groups mentioned in drug repurposing database are various neoplasms, immune system diseases, pathological signs and symptoms (clinical manifestations that can be either objective when observed by a physician, or subjective when perceived by the patient) and nervous system diseases. It is interesting to mention, that besides group C that comprises diseases, also repositioning indications can be in group E01 (Diagnosis), F02 (Psychological Phenomena and Processes), F03 (Mental Disorders), G08 (Reproductive and Urinary Physiological Phenomena), G11 (Musculoskeletal and Neural Physiological Phenomena). When checking the pairs of “original indication - secondary indication” for small molecule drugs (see Figure 2), in general, the most interesting cases are located in the area of intersection of C01 (bacterial infections), C02 and C03 (parasitic diseases) with other types of diseases. In such a case the repositioning can occur either to a homolog protein with conservation of the function or to a completely different protein target. In the first case a drug such as the antimycotic ketokonazole has been repositioned to a human target (Cytochrome P450) from its fungal homologus, to treat nephrotoxicity induced by cyclosporine. In the second case a drug such as doxycycline has been repurposed from bacterial infection as rpsD and rpsI inhibitor to stomatognathic disease as metalloproteinase inhibitor. However, this area is not particularly populated and cases of such repositioning are rare.

**Figure 2.**
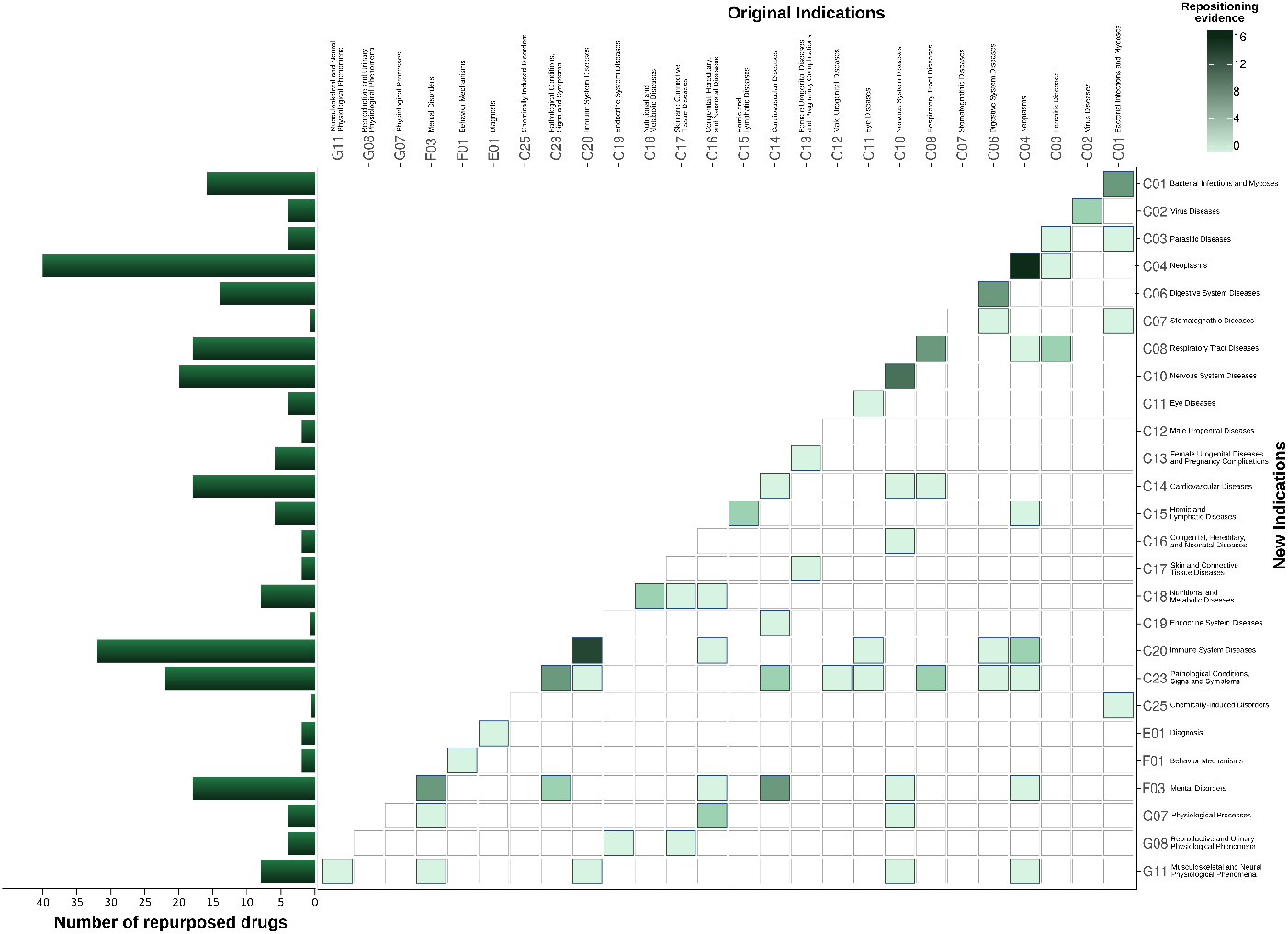
Frequency of disease among indication pairs. This figure shows the directions in terms of primary and secondary indications of all the repositioning cases analyzed. The disease classes have been plotted on both the axes and the number of repurposed drugs from one disease class to another has been expressed with the color intensity. The darkest squares lay on the central diagonal, showing that the majority of successful repositioning cases are carried within the same disease class. Besides on the Right side is displayed the number of repositioned drugs for the new indication.

When comparing the frequency of pairs “original indication - secondary indication” for diseases root MeSH key, as shown on the heatmap Figure 2 the main diagonal was the most populated, meaning 76 cases out of the 128 were repurposed within the same disease class, and thus belong to the Disease-centric group. The main cases are between one type of neoplasm against another type of neoplasm, cancer in the majority of cases, (C04 - C04), 16 cases in total, such as the kinase inhibitor Nilotinib, repurposed from Philadelphia chromosome positive chronic myelogenous leukemia to the treatment of gastrointestinal stromal tumors Table 1. Also very popular is repositioning for immune system diseases (C20 - C20), 13 cases in total, for which an example is the steroid beclomethasone repositioned from the treatment of rhinitis to the intestinal graft-versus-host disease Table 1; nervous system diseases (C10 - C10), 8 cases, such as midazolam HCL intravenous switched from preoperative sedation to epileptic seizure activity Table 1; and pathological conditions, signs and symptoms (C23 - C23), 5 cases, such as aminocaproic acid repurposed from enhancing hemostasis to topical treatment of traumatic hyphema of the eye Table 1. In reality, based on phenotypical similarities and handling approaches, even bigger group of brain related diseases and perception modification group C10-C23-F03 (mental disorders) exists. The popularity of main diagonal for drug repositioning can be explained by importance of the key protein targets for handling and treatment of many similar diseases at the same time.

**Table 1.**
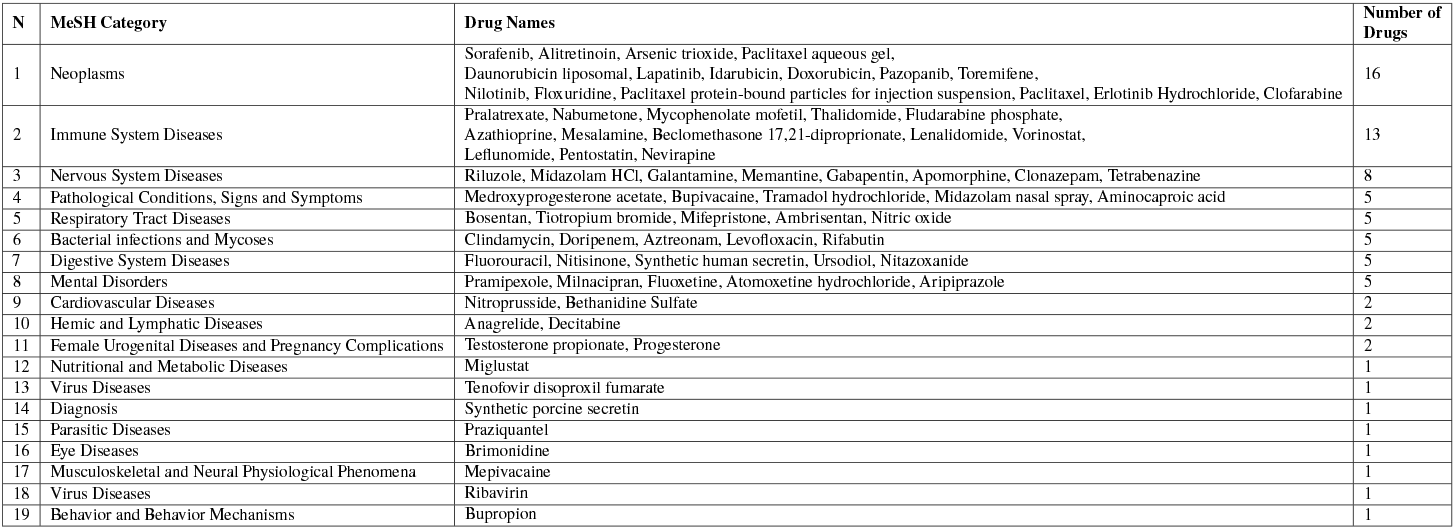
Disease-centric cases of repositioning. List of 76 disease-centric repositioned drugs grouped for indication category, according to the MeSH tree classification is shown. Since their original and secondary therapeutic indication falls within the same MeSH category no further analysis on the targets were carried.

### The 36% of the cases fall into the Target-centric category

The drug targets were linked to original and secondary indications using data mining and literature information. When the target was the same for both indications, or showed a sequence simialrity of at least 30%, the drug was classified as Target-centric. 6 identified cases were based on the binding of a drug to two homological targets with the same function (orthologue) (see Table 2). For ketokonazole amino-acid the similarity of targets was 49%, for eflornithine 57%, for dapsone 60%, for atovaquone 65% and for trimetrexate 66%. These values are much higher than the currently used 30 percent as a structural similarity threshold. In total, as shown in Table 2, 46 out of 128 drug cases were identified as Target-centric. Some examples include chlorpromazine, of which interaction with the serotonin receptor HTR2A is involved in both the antiemetic/antihistamine indication (Pathological Conditions, Signs and Symptoms) and the non-sedating tranquillizer action (Mental Disorder) Table 2; and celecoxib, non-steroideal antinflammatory, originally approved to treat osteoarthritis and adult rheumatoid arthritis (Immune System Diseases) for its interaction with the enzyme PTGS2, also known as COX-2, and subsequently repurposed to familial adenomatous polyposis (Congenital, Hereditary and Neonatal Diseases) with the same target Table 2.

**Table 2.**
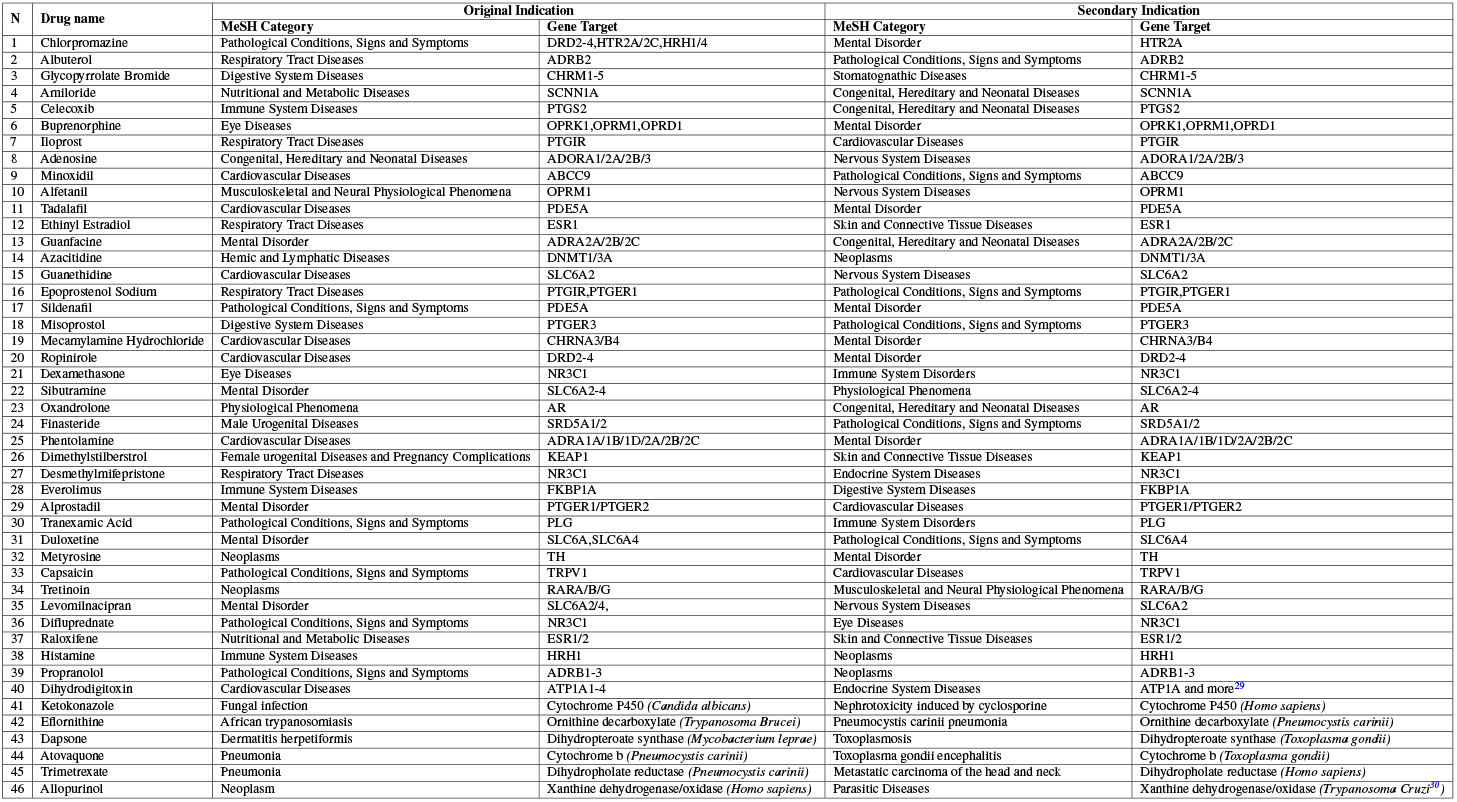
Taget-centric cases of repositioning. Disease (MeSH category) and protein targets (gene name or Uniprot ID) for both primary and secondary indications are shown. For the 46 cases of target-centric repositioning the target UniprotID is identical for both original and secondary indication. Target-disease associations retrieved from PubMed present a citation enclosed in the box. At the bottom, 6 drugs repurposed to an orthologue target.

### The 5% of the cases is classified as Drug-centric

The last step was made to identify cases of multi-target drugs. We searched for drugs for which the original and new indications were linked with different protein targets. Therefore, after completing the previous filtering steps, 6 out of 128 cases were left and classified as drug-centric. For example, valproic acid, used to treat episodes associated with bipolar disorder and seizures (nervous system diseases) by hitting the mithocondrial enzymes succinate-semialdehyde dehydrogenase (ALDH5A1) and 4-aminobutyrate aminotransferase (ABAT), repurposed to the treatment of familial adenomatous polyposis (congenital, hereditary and neonatal diseases) for the interaction with the histone deacetylase 2 (HDAC2) Table 3.

**Table 3.**
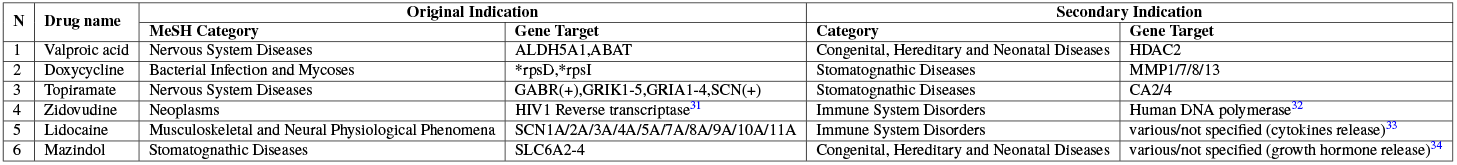
Drug-centric cases of repositioning. For each drug-centric case therapeutic indications (MeSH category) and protein targets (gene name or Uniprot ID (*:Non-human, (+):multiple subunits)) for both original and secondary indications are shown. According to our definition of drug-centric, those 6 cases must have a different MeSH code and protein target on the primary and secondary indication. Those 6 cases represents the most interesting situations in terms of diversity for the application of drug-target interaction prediction techniques. Target-disease associations retrieved from PubMed present a citation enclosed in the box.

## Discussion

### More than one third of the cases don’t fit the “small molecule drug - protein target” definition

Because of their higher promiscuous profiles and a more clear interaction behaviour with their targets, we exclusively focused on small molecule drugs. After the first step of filtering, a total of 68 cases out of the 196 were excluded, as they did not fit into “small molecule drug - protein target” scheme. For those cases the drug was usually an antibody and the target usually and RNA or another type of non-protein biomolecule. Examples of repositioned therapeutic antibodies were: infliximab, being used in Crohn’s disease and juvenile rheumatoid arthritis, adalimumab, also used in arthritis and Crohn’s disease. Therapeutic proteins were also present in the database, such as somatropin, that is being used to treat children with growth failure and for induction of ovulation in women with infertility. Some drugs had to be excluded because their action is linked to a non-protein target. For example, the DNA can be a direct target for some drugs, such as melphalan, used in patients with multiple myeloma and metastatic melanoma, or cladribine, used for treatment of hairy cell leukemia and chronic lymphocytic leukemia.

### Drug repositioning is mostly disease and target-centric

The retrospective analysis of the repositioning cases present in the RDD database gave us an interesting picture of the current state of drug repositioning. 60% of repositioned drugs analysed (76 cases out of 128) have been redirected to the same disease family. This tendency resulted to be extremely frequent within two categories of therapeutic indications: neoplasms and immune system disorders Figure 2, which show also the highest number of repositioned cases contained in our database. 30% of the analyzed drugs (46 out of 128) have been repurposed to a different condition but to the same protein target, intended as identical Uniprot ID. Only the 10% (6 cases) has been repositioned to a different disease and a different target. The situation hereby described seems to reflect a general trend common in the whole drug discovery world, which has been pointed as one of the reasons behind the structural crisis of the Pharmaceutical R&D mentioned in the introduction. Current pharmaceutical R&D situation has been compared to an drilling-oil process, where the cheapest and easiest opportunity with highest expected returns is exploit first and less attractive opportunities with lower returns are left behind^35^. Looking at our results, that could be translated in the prioritization of certain disease classes and fast repurposing approaches within the same disease and target families, with a big pool of drug-target-disease connections left almost entirely unexplored. In sight of this, it might be crucial in the future to invest in drug repositioning techniques focused on the study of fine charachteristics of drugs, targets and diseases (so called drug-centric approaches) and able to overcome the barrier represented by the permanence within the same disease and target category. A systematic and efficient repositioning approach able to exploit non-related diseases and targets might help both the pharmaceutical R&D by generating more profits and patients by bringing new therapeutic tools in a fast and cheap way.

### The role of drug-target interaction analysis in drug repositioning

Notwithstanding the computational drug repositioning has lately developed many strategies for drug-target interaction prediction, our analysis shows that most of the repositioning cases in our database might have not been obtained with those techniques. As shown in Figure 1, disease-centric drug repositioning can link a drug directly to a pathological condition with no need for assessing target similarity or analysing drug-target interactions. In that case no target-centric or drug-centric approach is necessary to obtain a new patent. For target-centric strategies firm link of target to indication is vital. Platforms such as open targets^36^ and Beagle^37^ are important to infer new connections, but often the link target-indication is not so direct and clear. The drug-centric cases recognized here are the only one who might have really benefit by drug-target interaction prediction methods (ligand-based, structure-based and machine learning-based).

### Limit and potential of Structure-Based drug repositioning as drug-centric approach

An example of drug-centric approach which aims to repurpose drugs to different targets and diverse indications is represented by the structure-based drug repositioning. This approach aims to predict new drug-target interactions by taking into account information about the structure of drugs and targets and their interactions. Although SBDR showed a big potential in repurposing known drugs to different targets and indications^25,38^, many limitations make this approach not so easy to apply in a relevant and systematic way. In fact, it requires the availability of structural data for both the original drug and the target as well as the putative new target with, possibly, its ligands. The lack of those information limits considerably the searching space for drug repositioning. In fact, for none of the cases classified in this work as drug-centric repositioning both the original drug-target couple and the repositioned complex were available on PDB, confirming the existing barriers in the application of SBDR.

### Inaccuracy in the drug repositioning process identification

It must be said that our analysis has been a retrospective work based on the final result of different repositioning processes and doesn’t account for the real process used to repurpose the drugs. Therefore we suspect that many drugs identified as result of disease-centric drug repositioning for the affinity between their first and second indication might be instead product of a target-centric or diseases-centric pipeline. For example, the drug nilotinib, which according our analysis falls in disease-centric drug repositioning, has been repurposed to a different protein target for gastrointestinal stromal tumor after target-affinity experiments^5^. For that reason, a deeper molecular understanding of drug-repositioning databases might lead to a more accurate analysis and give a clearer picture of the real status of repurposing.

## Methods

### Identification of current drug repositioning cases

The Repurposed Drug Database (RDD) was downloaded from www.drugrepurposingportal.com in January 2017. RDD contains 233 drugs that were repurposed up-to-date (see Annex II), providing information about the drugs with their original and new indications. Given the above, it was necessary to integrate this data with another source providing information about the protein targets involved in both indications.

### Retrieval of molecular drug targets data

A list of Molecular Drug Targets (MDT) was collected from the “Comprehensive map of molecular drug targets” provided by Santos et al.^39^, where authors curated “893 human and pathogen-derived biomolecules through which 1,578 US FDA-approved drugs act”. They used target annotations from the ChEMBL, DrugCentral and canSAR databases. The data was downloaded in .php format in January 2017 (see Annex II). The aim of that work was to facilitate mechanism-based drug discovery. The resulting drugs’ targets data set contained information about efficacy targets - targets to which the drug directly binds to exert its therapeutic effect (see Annex III). It is important to mention, that by definition of the work “biomolecules that the drug may also bind to, or be metabolized by, but which are not known to be responsible for its therapeutic effect, are not defined as targets”. Important to consider as well, is that the work defines drug as any therapeutic agent, including not only small molecules, but also other biological agents, that are used to enhance health.

### Identification of targets involved in the repositioning cases

Data in RDD were merged with data in MDT (drug targets) based on the commonly known drug name, resulting in 196 drugs matched by their common names, 263 unique targets and 333 unique indications (check Figure 3 or Annex I for more details). The merging of data was performed via a Python 2.7 script in April 2017 and complemented with other biological relevant databases in order to enrich the analysis (see Annex I).

**Figure 3.**
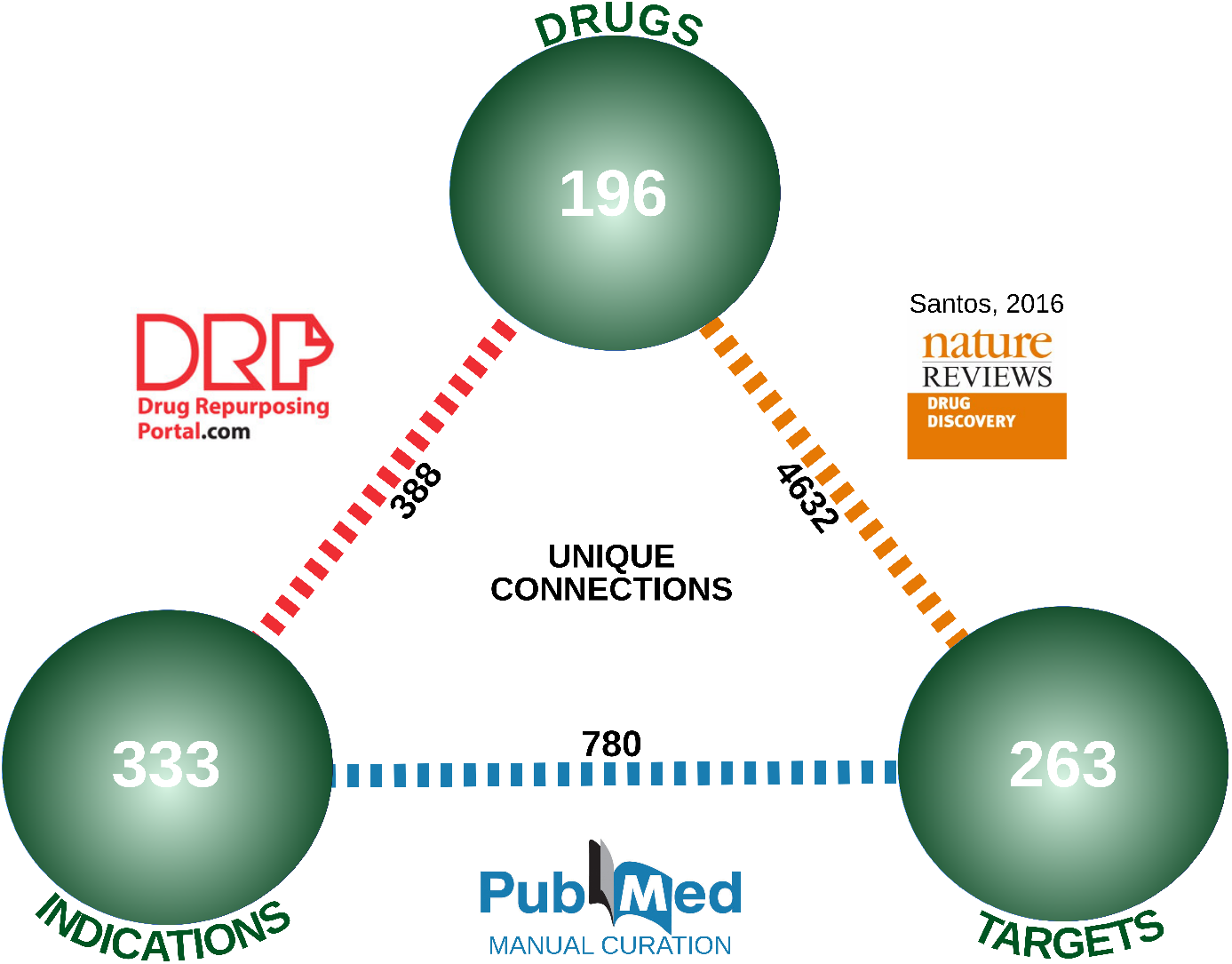
Collection and classification of known repositioning cases. Merging of Repositioned Drug Database (RDD), containing 196 drugs and 333 indications linked through 388 connections, the Molecular Target Database from Santos (MTD), containing 4632 links between 196 drugs and 263 targets, and PubMed, which allowed to find 780 different links between 263 targets and 333 indications.

### Filter of not small molecules and not protein targets

This study only considered proteins as biomolecule targets and small molecule as drugs. Other cases, such as antibodies as drugs and other biomolecules (e.g enzymes or unknown) than proteins as targets, were left outside. Out of the 196 drugs both in RDD and MDT, a total of 68 were removed under this filtering criteria (see Annex II).

### Identification of Disease-centric repositioning cases

A repositioning case is considered as Disease-centric when it is based on the similarity of phenotype. For each drug case, the MeSH tree root key was assigned to each indication (see Table 4). The frequency of diseases among repurposed cases was calculated and visualized with the Matplotlib python library and the number of cases for each pair of root MeSH keys were plotted in R with the ggplot2 package (see Figure 2). For each pair, all targets mentioned for the cases at the intersection of the same root MeSH keys were collected and the number of targets at the intersection calculated. Literature evidence about applicability of these targets for the group was also collected.

**Table 4.** Root MeSH tree key for the group of diseases and corresponding name. MeSH category codes (left column) and common names (right column) for all types of diseases and conditions for which drug repositioning was previously applied according to RDD.

| MeSH Category | Disease group | MeSH Category | Disease group |
| --- | --- | --- | --- |
| C01 | Bacterial Infections and Mycoses | C16 | Congenital, Hereditary, and Neonatal Diseases and Abnormalities |
| C02 | Virus Diseases | C17 | Skin and Connective Tissue Diseases |
| C03 | Parasitic Diseases | C18 | Nutritional and Metabolic Diseases |
| C04 | Neoplasms | C19 | Endocrine System Diseases |
| C06 | Digestive System Diseases | C20 | Immune System Diseases |
| C07 | Stomatognathic Diseases | C23 | Pathological Conditions, Signs and Symptoms |
| C08 | Respiratory Tract Diseases | C25 | Chemically-Induced Disorders |
| C10 | Nervous System Diseases | E01 | Diagnosis |
| C11 | Eye Diseases | F01 | Behaviour Mechanisms |
| C12 | Male Urogenital Diseases | F03 | Mental Disorders |
| C13 | Female Urogenital Diseases and Pregnancy Complications | G07 | Physiological processes |
| C14 | Cardiovascular Diseases | G08 | Reproductive and Urinary Physiological Phenomena |
| C15 | Hemic and Lymphatic Diseases | G11 | Musculoskeletal and Neural Physiological Phenomena |

### Target assignment to original and secondary indication

In order to distribute targets into original and secondary indications we used literature evidence in PubMed (see Figure 3). Target-indication connections were retrieved manually from literature by searching for direct evidences, such as the improvement in a condition upon treatment with a certain drug due to action on the protein target; indirect evidences, in case the correlation between a condition with a certain target activity and a disease were reported by two different articles; or generalized evidences, if a small single evidence on target-condition link was found even though no strong correlation has been yet recognized. The validation for the previous manual-curation was done via text-mining by using the ensemble biclustering algorithm (EBC), which allow to extract the connections from a natural text in a machine-processable form 42. The text mining dataset used in this work consists of 2 parts. Part I connects dependency paths to labels, or ‘themes’. They were introduced in this dataset to label what kind of interaction exist between two terms, e.g. whether a casual mutation has a role in pathogenesis or promotes progression of the disease. The second part of the dataset contains information about drug-taret-indication associations. To validate manual PubMed curation and estimate how applicable is text-mining for this aim in general, target-indication associations in this dataset were used. To make it possible to use text-mining dataset with the drug repositioning dataset with targets, UniProt IDs were turned into gene IDs using UniProt API service, and MeSH on demand was used to assign indications IDs to textual description of diseases. Afterwards, all the records, containing the genes of a proteins assigned as a therapeutic targets for drug repositioning, were collected from the dataset, and checked whether the pair gene ID - MeSH ID is present. To identify, whether the gene encodes a drug target, the Part I and Part II of text mining dataset were linked and selected entries were the gene actually had ‘drug target’ label. In this way targets were again distributed into original and secondary indication (see Annex III). The resulting distribution was compared to manual distribution.

### Identification of Target-centric repositioning cases

A repositioning case is considered as Target-centric when exploits the same protein target but in different cellular contexts. The remaining cases which were not classified as Disease-centric repositioning, were analyzed in order to determine whether the drug actually has two different indications related to the same target or not. Therefore, cases for which UniProt IDs were the same, were marked as target-centric repositionings.

### Identification of Drug-centric repositioning cases

A repositioning case is considered as Drug-centric when exploits drug’s chemical properties. First, the cases which were not classified as Disease-centric and neither Target-centric were considered as potential Drug-centric. Cases for which there was a literature evidence that targets are different, but secondary target was not mentioned by the comprehensive map were marked as Drug-centric as well. Even though, some targets have different Uniprot IDs they can be homological targets with the same function in different organisms (ortholog). In order to evaluate the above, targets sequence similarity was evaluated using Clustal Omega alignment (https://www.ebi.ac.uk/Tools/msa/clustalo) where the number of similar and identical positions was summarized and divided be the length of the alignment (usually length of the longest sequence).

### Drug-centric cases with structure-based drug repositioning

We finally evaluated how feasible is for the structure-based drug repositioning approach to identify the selected Drug-centric cases as repositioning stories. To do so, a mapping file from Uniprot to PDB ID was generated and used to search all available structures in Protein Data Bank 43, describing the binding between the drugs and their corresponding targets (associated to both indications). In addition to the previous step, structures describing a binding between the drugs but with another target (not described in the MDT) were also considered and evaluated under the terms of structure-based drug repositioning.

## Conclusion

Drug-target interaction prediction is an important part of most of the rational drug repositioning pipelines. In fact, different biochemical, physical and mathematical techniques have been designed and optimized to infer accurately links between ligands and proteins. In this work we analyzed many successful drug-repositioning cases and, based on the similarity between old and new indication and old and new target, we evaluated the real impact of drug-target interaction prediction for them. Dividing all the cases containing small molecules and protein targets (128) in Disease-centric (with very similar indications), Target-centric (with identical or orthologue targets) and Drug-centric cases (with different targets in different indications), we concluded that only 6 cases out of 128 would have needed drug-target interaction prediction to rationally infer the repurposing. This unexpected small number of drugs repurposed to a different target in a different indication might be due to the high costs in terms of information, time and money required by drug-target interaction prediction approaches compared to Target- and Disease-centric ones. On the other hand, those results highlight a big unexplored niche for drug-target interaction prediction in drug repositioning, which might be safely explored with the ongoing advancements of techniques designed to infer a link between a ligand and a new protein target.

## Supporting information

Supplemental Table III, list of target-indication-drug links

Supplemental Table II, list of RDD repurposed drugs

Supplemental Table I, resulting merged cases of repurposed drugs

